# Non-invasive detection of bladder cancer through the analysis of driver gene mutations and aneuploidy

**DOI:** 10.1101/203976

**Authors:** Simeon Springer, Maria Del Carmen Rodriguez Pena, Lu Li, Christopher Douville, Yuxuan Wang, Josh Cohen, Diana Taheri, Bahman Afsari, Natalie Silliman, Joy Schaeffer, Janine Ptak, Lisa Dobbyn, Maria Papoli, Isaac Kinde, Bahman Afsari, Aline C. Tregnago, Stephania M. Bezerra, Christopher VandenBussche, Kazutoshi Fujita, Dilek Ertoy, Isabela W. Cunha, Lijia Yu, Mark Schoenberg, Trinity J. Bivalacqua, Kathleen G. Dickman, Arthur P. Grollman, Luis A. Diaz, Rachel Karchin, Ralph Hruban, Cristian Tomasetti, Nickolas Papadopoulos, Kenneth W. Kinzler, Bert Vogelstein, George J. Netto

## Abstract

Current non-invasive approaches for bladder cancer (BC) detection are suboptimal. We report the development of non-invasive molecular test for BC using DNA recovered from cells shed into urine. This “UroSEEK” test incorporates assays for mutations in 11 genes and copy number changes on 39 chromosome arms. We first evaluated 570 urine samples from patients at risk for BC (microscopic hematuria or dysuria). UroSEEK was positive in 83% of patients that developed BC, but in only 7% of patients who did not develop BC. Combined with cytology, 95% of patients that developed BC were positive. We then evaluated 322 urine samples from patients soon after their BCs had been surgically resected. UroSEEK detected abnormalities in 66% of the urine samples from these patients, sometimes up to 4 years prior to clinical evidence of residual neoplasia, while cytology was positive in only 25% of such urine samples. The advantages of UroSEEK over cytology were particularly evident in low-grade tumors, wherein cytology detected none while UroSEEK detected 67% of 49 cases. These results establish the foundation for a new, non-invasive approach to the detection of BC in patients at risk for initial or recurrent disease.

## Introduction

Bladder cancer (BC) is the most common malignancy of the urinary tract. According to the American Cancer Society, 79,030 new cases of bladder cancer and 18,540 deaths are estimated to occur in the United States alone in 2017 [1]. Predominantly of urothelial histology, invasive BC arises from non-invasive papillary or flat precursors. Many BC patients suffer with multiple relapses prior to progression, providing ample lead-time for early detection and treatment prior to metastasis [2]. Urine cytology and cystoscopy with transurethral biopsy (TURB) are currently the gold standards for diagnosis and follow-up in bladder cancer. While urine cytology has value for the detection of high-grade neoplasms, it is unable to detect the vast majority of low-grade tumors [3-5]. This fact, together with the high cost and invasive nature of repeated cystoscopy and TURB procedures, have led to many attempts to develop novel noninvasive strategies. These include urine or serum based genetic and protein assays for screening and surveillance [6-21]. Currently available U.S. Food and Drug Administration (FDA) approved assays include ImmunoCyt test (Scimedx Corp), nuclear matrix protein 22 (NMP22) immunoassay test (Matritech), and multitarget FISH (UroVysion) [6-12]. Sensitivities between 62% and 69% and specificities between 79% and 89% have been reported for some of these tests. However, due to assay performance inconsistencies, cost or required technical expertise, integration of such assays into routine clinical practice has not yet occurred.

Much is now known about the genetic pathogenesis of BC. High rates of activating mutations in the upstream promoter of the *TERT* gene are found in the majority of BC as well as in other cancer types [22-24]. *TERT* promoter mutations predominantly affect two hot spots, g.1295228 C>T and g.1295250 C>T. They lead to the generation of CCGGAA/T or GGAA/T motifs altering binding site for ETS transcription factors and subsequently increased *TERT* promoter activity [22, 25]. *TERT* promoter mutations occur in up to 80% of invasive urothelial carcinomas of the bladder and upper urinary tract as well as in several of its histologic variants [14, 23, 26-28]. Moreover, *TERT* promoter mutations occur in 94 60-80% of BC precursors, including Papillary Urothelia Neoplasms of Low Malignant Potential [29], non-invasive Low Grade Papillary Urothelial Carcinoma, non-invasive High Grade Papillary Urothelial Carcinoma and “flat” Carcinoma in Situ (CIS), as well as in urinary cells from a subset of these patients [14]. *TERT* promoter mutations have thus been established as the most common genetic alteration in BC [14, 30]. Other important oncogene-activating mutations include those in *FGFR3*, *RAS* and *PIK3CA*, which have been shown to occur in a high fraction of non-muscle invasive bladder cancers [31, 32]. In muscle-invasive bladder cancers, mutations in *TP53*, *CDKN2A*, *MLL2* and *ERBB2* are also frequently found [33-40]

The current study assesses the performance of a massively parallel sequencing-based assay, termed UroSEEK, for the detection of BC through the analysis of urinary cells. UroSEEK has three components: detection of intragenic mutations in regions of ten genes (*FGFR3*, *TP53*, *CDKN2A*, *ERBB2*, *HRAS*, *KRAS*, *PIK3CA*, *MET*, *VHL and MLL2*) that are frequently mutated in BC [33-40]; detection of mutations in the *TERT* promoter [22-24]; and detection of aneuploidy [41, 42]. UroSEEK was applied to two independent cohorts of patients. The first (called the Early Detection cohort) involved patients with microscopic hematuria or dysuria, which are both risk factors for BC. Only a relatively small fraction (4 to 5%) of micro-hematuria patients are at risk for developing urothelial malignancy [43, 44], so the decision about which of these patients should undergo cystoscopy is often difficult. The second cohort (called the Surveillance cohort) involved patients who had already been diagnosed with BC. These patients are at highn risk for recurrence [43]. Because urine cytology is relatively insensitive for the detection of recurrence, cystoscopies are performed as often as every three months in such patients in the U.S. In fact, the cost of managing these patients is in aggregate higher than the cost of managing any other type of cancer, and amounts to 3 billion dollars annually [45]. A non-invasive test that could predict which of these patients were most likely to develop recurrent BC could thereby be both medically and economically important.

## Results

A schematic of the approach used in this study is provided in Figure 1A, and a flow diagram indicating the number of patients evaluated in this study and the major results is provided in Figure 1B.

**Figurre 1A.**
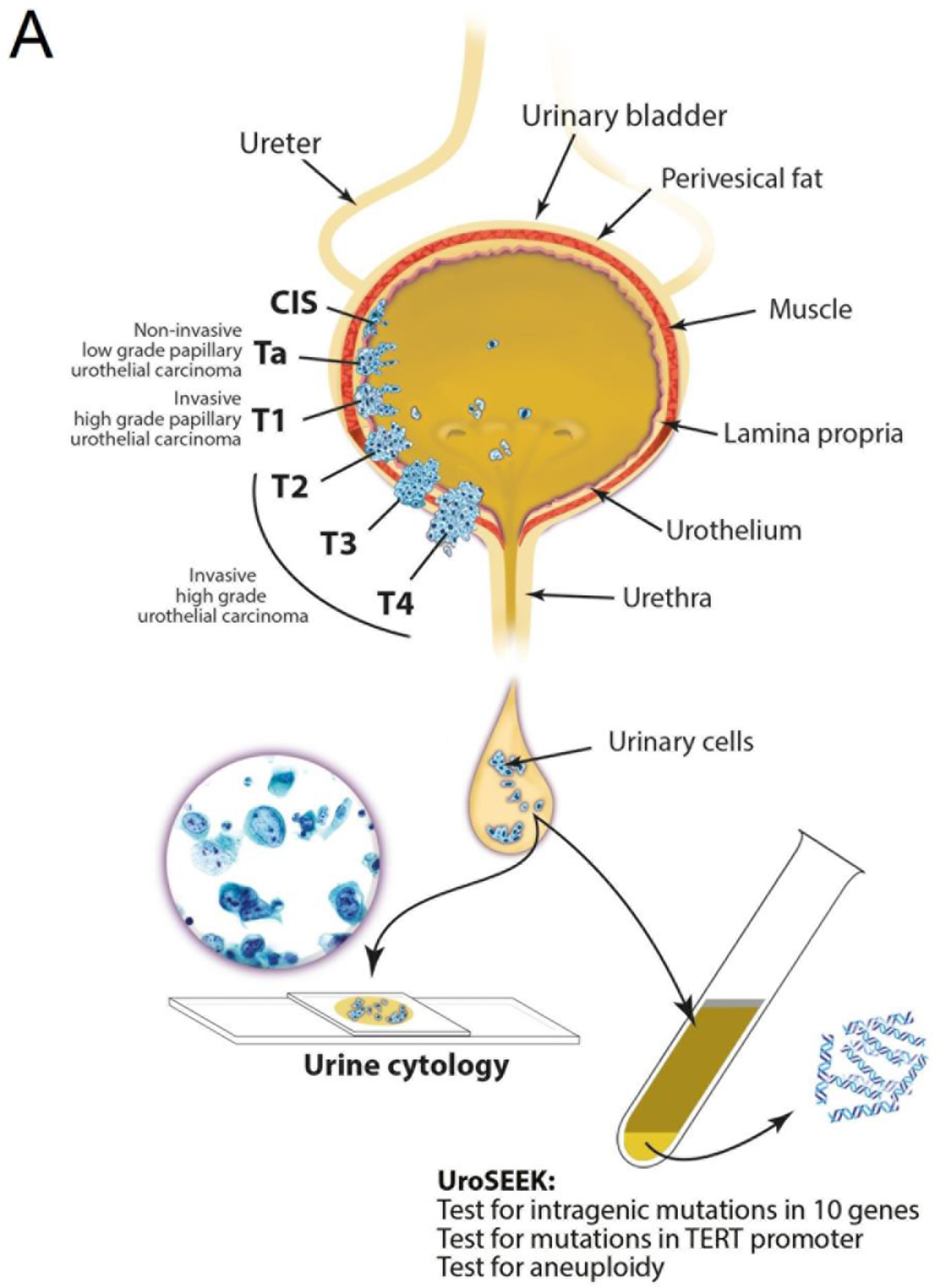
Schematic of the approach used to evaluate urinary cells in this study.

**Figure 1B.**
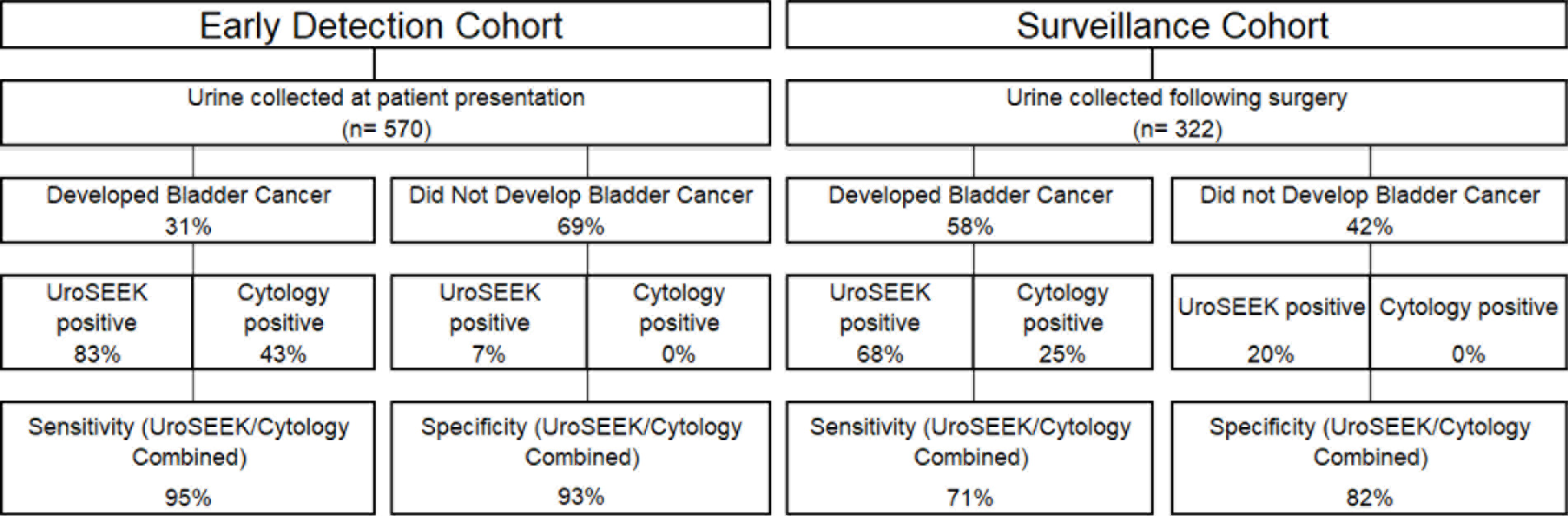
Flow diagram indicating the number of patients in the two cohorts evaluated in this study and summarizing the salient findings. Cytology was performed on only a subset of the patients (see main text).

## Early Detection Cohort

### Cohort characteristics

A total 570 patients were included in the Early Detection cohort, each with one urine sample analyzed. 90% of the patients had hematuria, 3% had lower urinary tract symptoms (LUTS), and 9% had other indications suggesting they were at risk for BC. The median age of the participants was 58 years (range 5 to 89) (Supplementary Table 1). As expected from prior studies of patients at risk for BC, 70% of the patients were male [1, 43]. 175 (31%) of patients developed BC after a median follow-up period of 18 months (range 0 to 40 months). For each patient who developed BC, we selected two other patients who presented with similar symptoms but did not develop BC during the follow-up period. By design, then, the fraction of cases in this cohort developing BC was higher than the fraction (5%) of patients with similar presentations that would have developed BC in standard clinical practice. The characteristics of the tumors developing in the 570 patients are summarized in Supplementary Table 1 and detailed in Supplementary File 1.

### Genetic analysis

We performed three separate tests for genetic abnormalities that might be found in urinary cells derived from BC (Figure 2). First, we evaluated mutations in selected regions of ten genes that have been shown to be frequently altered in urothelial tumors (Supplementary File 2). For this purpose, we designed a specific set of primers that allowed us to detect mutations in as few as 0.03% of urinary cells. The capacity to detect such low mutant fractions was a result of the incorporation of molecular barcodes in each of the primers, thereby substantially reducing the artifacts associated with massively parallel sequencing [41]. Second, we evaluated *TERT* promoter mutations. A singleplex PCR was used for this analysis because the unusually high GC-content of the *TERT* promoter precluded its inclusion in the multiplex PCR design. Third, we evaluated the extent of aneuploidy using a technique in which a single PCR is used to co-amplify ~38,000 members of a subfamily of long interspersed nucleotide element-1 (L1 retrotransposons, also called LINEs). L1 retrotransposons, like other human repeats, have spread throughout the genome via retrotransposition and are found on all 39 non-acrocentric autosomal arms [41].

**Figure 2.**
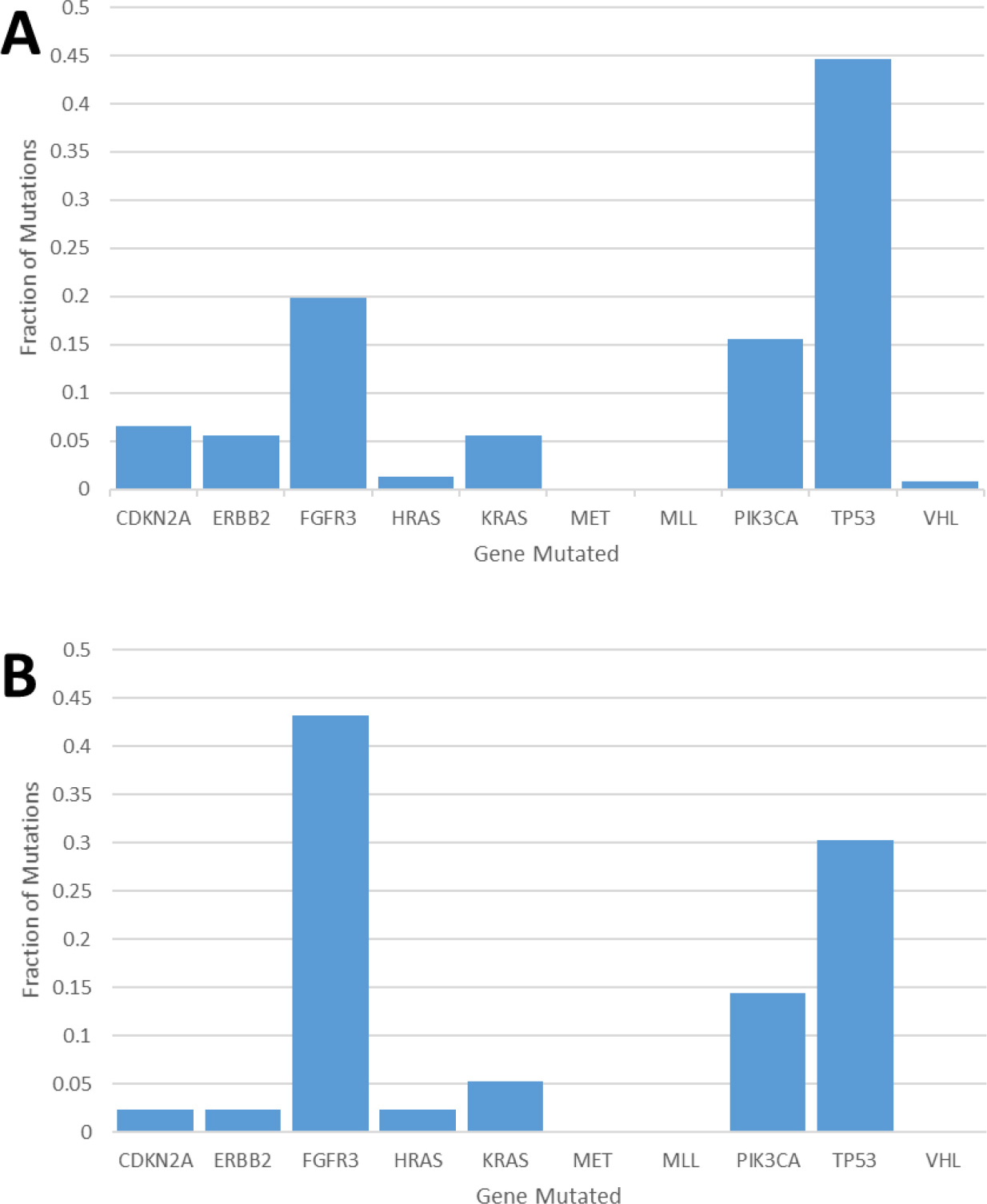
Fraction of mutations found in the ten-gene panel in 231 urinary cell samples assessed in the (A) early detection cohort and 132 urinary cell samples assessed in the (B) surveillance cohort.

The multiplex assay detected mutations in 68% of the 175 urinary cell samples from the individuals that developed BC during the course of this study (95% CI 61% to 75%) (Supplementary Table 1 and Supplementary File 2). A total of 246 mutations were detected in 8 of the ten target genes (Figure 2A and Supplementary File 2). The mean mutant allele frequency in the urinary cells with detectable mutations was 18% and ranged from 0.17% to 99%. The most commonly altered genes were TP53 (45% of the total mutations) and FGFR3 (20% of the total mutations; Fig. 2A). The distribution of mutant genes was roughly consistent with expectations based on previous exome-wide sequencing studies of BCs [46]. At the thresholds used, 1.7% of the 395 patients in the Early Detection Cohort who did not develop BC during the course of the study had a detectable mutation in any of the ten genes. At the same thresholds, none of the 188 urinary cell samples from healthy individuals had any mutation in any of the ten genes assayed (100% specificity, 95% CI 98% to 100%).

Mutations in the *TERT* promoter were detected in 57% of the 175 urinary cell samples from the patients that developed cancer during the study interval (95% Ci 49% to 64%; Supplementary Table 1 and Supplementary File 3). The mean *TERT* mutant allele frequency in the urinary cells was 14% and ranged from 0.18% to 78%. Mutations were detected in 3 positions: 98% of the mutations were at hg1295228 (79%) and hg 1295250 (19%), which are 66 and 88 bp upstream of the transcription start site, respectively. These positions have been previously shown to be critical for the appropriate transcriptional regulation of *TERT*. In particular, the mutant alleles recruit the GABPA/B1 transcription factor, resulting in the H3K4me2/3 mark of active chromatin and reversing the epigenetic silencing present in normal cells [47]. 4% of the 395 patients in this cohort who did not develop BC during the course of the study had a detectable mutation in the *TERT* promoter. Only one of the 188 urinary samples from healthy individuals harbored a *TERT* promoter mutation.

Aneuploidy was detected in 46% (95% CI 39% to 54%) of the 175 urinary cell samples from the patients that developed BC during the course of the study, Supplementary Table1 and Supplementary File 4. The most commonly altered arms were 5q, 8q, and 9p. All three of these arms harbor well-known oncogenes and tumor suppressor genes which have been shown to undergo copy number alterations in many cancers, including BC [48]. 1.5% of the urinary cell samples from the 395 patients who did not develop BC during the course of the study exhibited aneuploidy. None of the 188 urinary samples from healthy individuals exhibited aneuploidy when assessed with the same technology.

### Comparison with primary tumors

Tumor samples from 102 of the patients enrolled in this cohort were available for comparison and were studied with the same three assays used to study the urinary cell samples (Supplementary File 3). In 91 (89%) of these 102 cancers, at least one mutation in the eleven genes studied were mutated (in the 10-gene panel or in the *TERT* promoter). Moreover, at least one of the mutations identified in the urine samples from these 102 patients was also identified in 83% of the corresponding BC (Supplementary Files 2 and 3). Analysis of the BCs also shed light on the basis for “false negatives”, i.e., the reason that 21% of urine samples from patients who developed BC had no detectable mutations in the 11 genes tested. The reason could either have been that the corresponding BC did not harbor a mutation in these 11 genes or that it did, but the fraction of neoplastic cells in the urine sample was not high enough to allow its detection with the assays we used. We could identify a mutation in at least one of the 11 genes in 62% of the primary tumors from patients with false negative urine tests for mutations (Supplementary File 5 and 6). We conclude that 38% of the 29 false negative tests for mutations were due to the fact that none of the queried mutations were present in the tumor and that the other 62% of the false negatives were due to insufficient amounts of cancer cells in the urine.

### UroSEEK: biomarkers in combination

As noted above, the ten-gene multiplex assay, the *TERT* singleplex assay, and the aneuploidy assays yielded 68%, 57%, and 46% sensitivities, respectively, when used separately (Supplementary Table 1 and Supplementary Files 2, 3, and 4). 45 samples without *TERT* promoter mutations could be detected by mutations in one of the other ten genes (Figure 3A and Supplementary File 2). Conversely, 35 samples without detectable mutations in the multiplex assay could be detected by virtue of *TERT* promoter mutations (Figure 3A and Supplementary File 3). Ten of the urinary cell samples without any detectable mutations in the 11 genes could be detected by the assay for aneuploidy (Figure 3A and Supplementary Table 4). Thus, when the three assays were used together (test termed “UroSEEK“), and a positive result in either assay was sufficient to score a sample as positive, the sensitivity rose to 83% (95% CI 76% to 88 %). Only one of the 188 samples from healthy individuals was scored positive by UroSEEK (specificity 99.5%, CI 97% to 100%). Twenty-six (6.5%) of the 395 patients in this cohort who did not develop BC during the course of the study scored positive by the UroSEEK test (specificity 93%, CI 91% to 96%). On average, UroSEEK positivity preceded the diagnosis of BC by 2.3 months, and in eight cases by more than a year (Figure 4A and Supplementary File 1).

**Figure 3.**
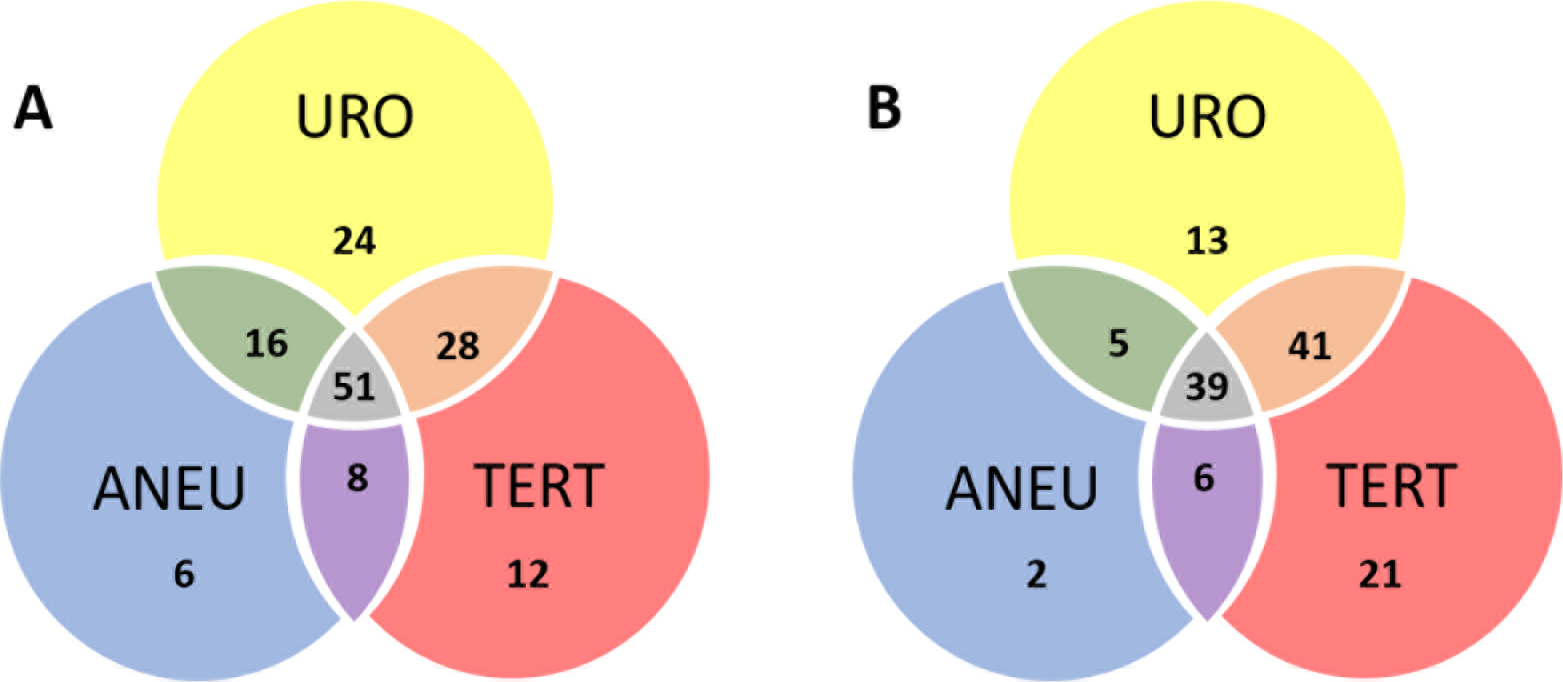
Venn Diagram of the distribution of samples that were positive by each of the three assays for the (A) early detection and (B) surveillance cohorts. URO = Ten gene panel, *TERT* = *TERT* promoter region, ANEU = Aneuploidy test.

**Figure 4.**
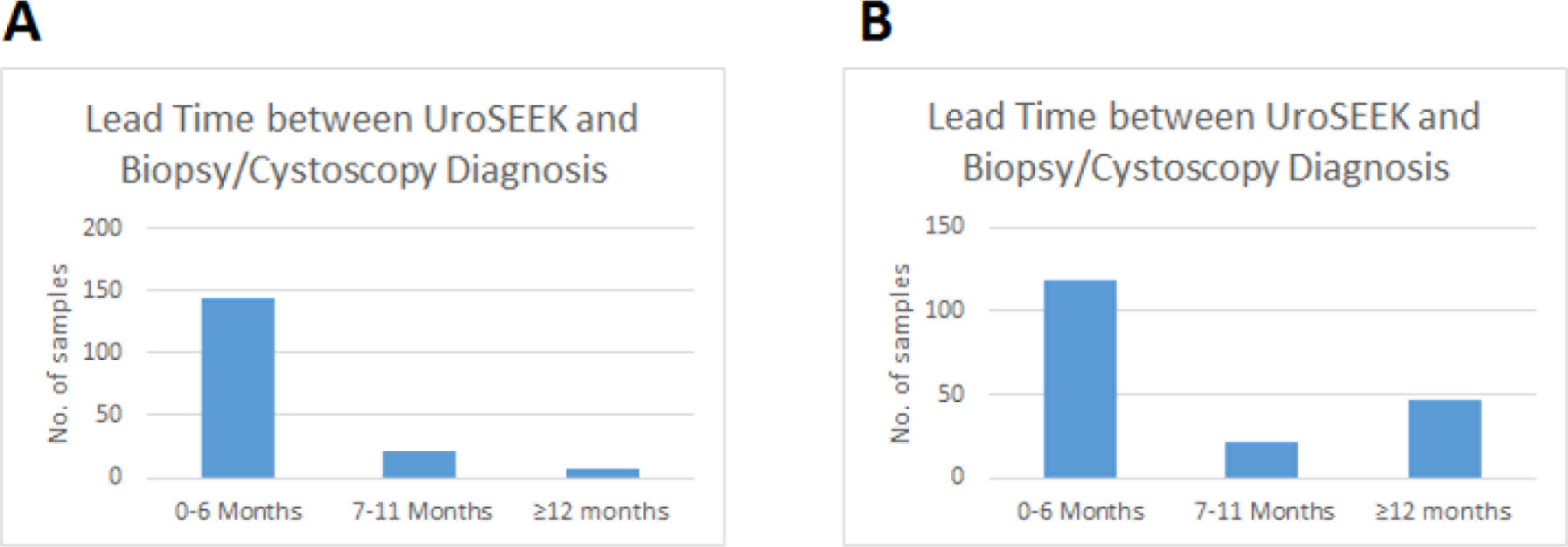
Bar graphs of the lead time between a positive UroSEEK test and the detection of disease at the clinical level in the (A) Early Detection and (B) Surveillance cohorts.

### UroSEEK plus cytology

As both cytology and UroSEEK tests are non-invasive and can be performed on the same urine sample, we assessed their performance in combination. There were 347 patients in the Early Detection cohort in whom cytology was available (Supplementary Table 1 and Supplementary File 1). Among the 40 patients who developed biopsy-proven cancer in this cohort, 17 were positive by cytology (43% sensitivity). None of the 299 patients that did not develop cancer were positive by cytology (100% specificity). UroSEEK was positive in 100% of the 17 cancer patients whose urines were positive by cytology and in 95% of the 23 cancer patients whose urines were negative by cytology. Thus, in combination, UroSEEK plus cytology afforded 95% (95% CI 83% to 99%) sensitivity, a 12% increase over UroSEEK and a 52% increase over cytology. Among the 299 patients in the early detection cohort who did not develop BC during the course of the study, 20 (6.6%) were positive by UroSEEK or cytology, giving the combination of UroSEEK and cytology a specificity of 93% (95% CI 90% to 96%).

## Surveillance Cohort

### Cohort characteristics

Our strategy for surveillance was different than the one we used for early detection. Patients in whom a BC was surgically excised for treatment and diagnosis generally have tumor tissue available, and in most such tumors, a mutation can be identified. For example, we found during the course of this study that a mutation in at least one of the 11 queried genes was present in 95.2% of BCs evaluated. All patients selected for the surveillance study had biopsy confirmed BC and had a urine sample collected 0 - 5 years after surgery. We were to evaluate a total of 322 patients that donated urines and whose BC contained a mutation in at least one of the 11 genes evaluated. We determined whether a single urine sample taken a relatively short time following surgical excision of the BC could reveal residual disease in these 322 patients, as evidenced by later recurrence. 187 (58%) of the 322 patients developed clinically evident BC after a median follow-up period of 10.7 months (range 0 to 51 months). The histopathologic types and tumor stages of these patients are summarized in Supplementary Table 2 and detailed Supplementary File 7. The median age of the participants was 62 (range 20 to 93). As expected from the demographics of BC, 75% of the patients were male.

### Genetic analysis

The multiplex assay in urinary cells detected mutations in 49% of the urinary cell samples from patients that developed recurrent BC during the study interval (95% CI 45% to 60%; Supplementary File 7 and Supplementary File 8). The mean mutant allele frequency in the urinary cells with detectable mutations was 16% and ranged from 0.08% to 93%. The most commonly altered genes were FGFR3 (43% of the 134 mutations) and TP53 (30% of the 134 mutations; Figure 2B). Seven percent of the 135 patients who did not develop recurrent BC during the course of the study had a detectable mutation in their urinary cell sample (these are *considered* to be false positives; see Discussion). The mean interval between a positive multiplex assay test and the diagnosis of recurrent BC was 7 months (range 0 to 51 months).

Mutations in the *TERT* promoter were detected in 51% of the urinary cell samples from patients that developed recurrent BC during the study interval (95% CI 44% to 58%; Supplementary Table 2 and Supplementary File 9). The mean *TERT* mutant allele frequency in the urinary cells with detectable mutations was 6% and ranged from 0.23% to 43%. Mutations were detected in the same three positions observed in the urinary cells of the Early Detection cohort. 10% (95% CI 83% to 94%) of the 135 patients who did not develop recurrent BC during the course of the study had a detectable *TERT* promoter mutation in their urine sample (false positives). The mean interval between a positive *TERT* test and the diagnosis of recurrent BC was 7 months (range 0 to 40 months).

Aneuploidy was detected in 30% (95% CI 24% to 37%) of the urinary cell samples from the patients that developed recurrent BC during the course of the study (Supplementary Table 2 and Supplementary File 10). The most commonly altered arms were 8p, 8q, and 9p, as in the Early Detection cohort. Two percent of the 135 patients who did not develop recurrent BC during the course of the study exhibited aneuploidy in at least one of their urinary cell samples.

### Markers in combination

As noted above, the ten-gene multiplex assay, the *TERT* singleplex assay, and the aneuploidy assays yielded 49%, 51%, and 30% sensitivities, respectively, when used separately (Supplementary Table 2 and Supplementary Files 8, 9 and 10). Thirty-two samples without *TERT* promoter mutations could be detected by mutations in one of the other ten genes (Figure 3B and Supplementary File 8). Conversely, 41 samples without detectable mutations in the multiplex assay could be detected by virtue of *TERT* promoter mutations. Three of the urinary cell samples without any detectable mutations could be detected by the assay for aneuploidy. Thus, the sensitivity of UroSEEK was 66% (95% CI 59% to 73%) Supplementary Table 2). Fouteen percent of the 135 patients in this cohort who did not develop BC during the course of the study scored positive by the UroSEEK test, yielding a specificity of 86% (95% CI 77% to 91%). On average, UroSEEK positivity preceded the diagnosis of BC by 7 months, and in 47 cases by more than one year (Figure 4B and Supplementary File 7).

There were 196 patients in the Surveillance cohort in whom cytology was available (Supplementary File 7). Among the 120 patients who developed recurrent BC in this cohort, 30 (25%) were positive by cytology. Conversely, no positive cytology results were observed in patients whose tumors did not recur. UroSEEK was positive in 90% of the recurrent BC patients whose urines were positive by cytology and in 61% of the 90 recurrent BC patients whose urines were negative by cytology. Thus in combination, UroSEEK plus cytology afforded 71% sensitivity (95% CI 61.84% to 78.77) (Figure 3D and Supplementary File 5). Among the 76 patients who did not develop recurrent BC during the course of the study and in whom cytology was available, 18% scored as positive by either cytology or UroSEEK, affording a specificity of 82% (95% CI 71% to 90%; see Discussion).

## Low vs. high grade urothelial neoplasms (both cohorts)

The advantage of UroSEEK over cytology was particularly evident in low-grade tumors (Papillary urothelial neoplasms of low malignant potential and non-invasive low grade papillary urothelial carcinomas). There were a total of 49 low-grade tumors evaluated in this study in whom cytology was available (six from the Early Detection cohort and 43 from the Surveillance cohort). None of these low-grade tumors were detected by cytology (0% sensitivity; 95% CI 0.0% to 6.7%) In contrast, UroSEEK detected 67% (95% CI 51% to 81%) of the low-grade tumors (identical rate of 67% in both cohorts; Supplementary Table 3 and Figure 5). Analogously, there were a total of 102 high-grade tumors (in-situ urothelial carcinoma, non-invasive high grade papillary urothelial carcinoma or infiltrating high grade urothelial carcinoma) evaluated in this study in whom cytology was available (34 from the Early Detection cohort and 68 in the Surveillance cohort). Cytology was positive in 45% of these patients (50% and 41 % in the Early Detection and Surveillance cohorts, respectively) while UroSEEK was positive in 80% of them (100% and 71% in the Early Detection and Surveillance cohorts, respectively; Supplementary Table 3).

**Figure 5.**
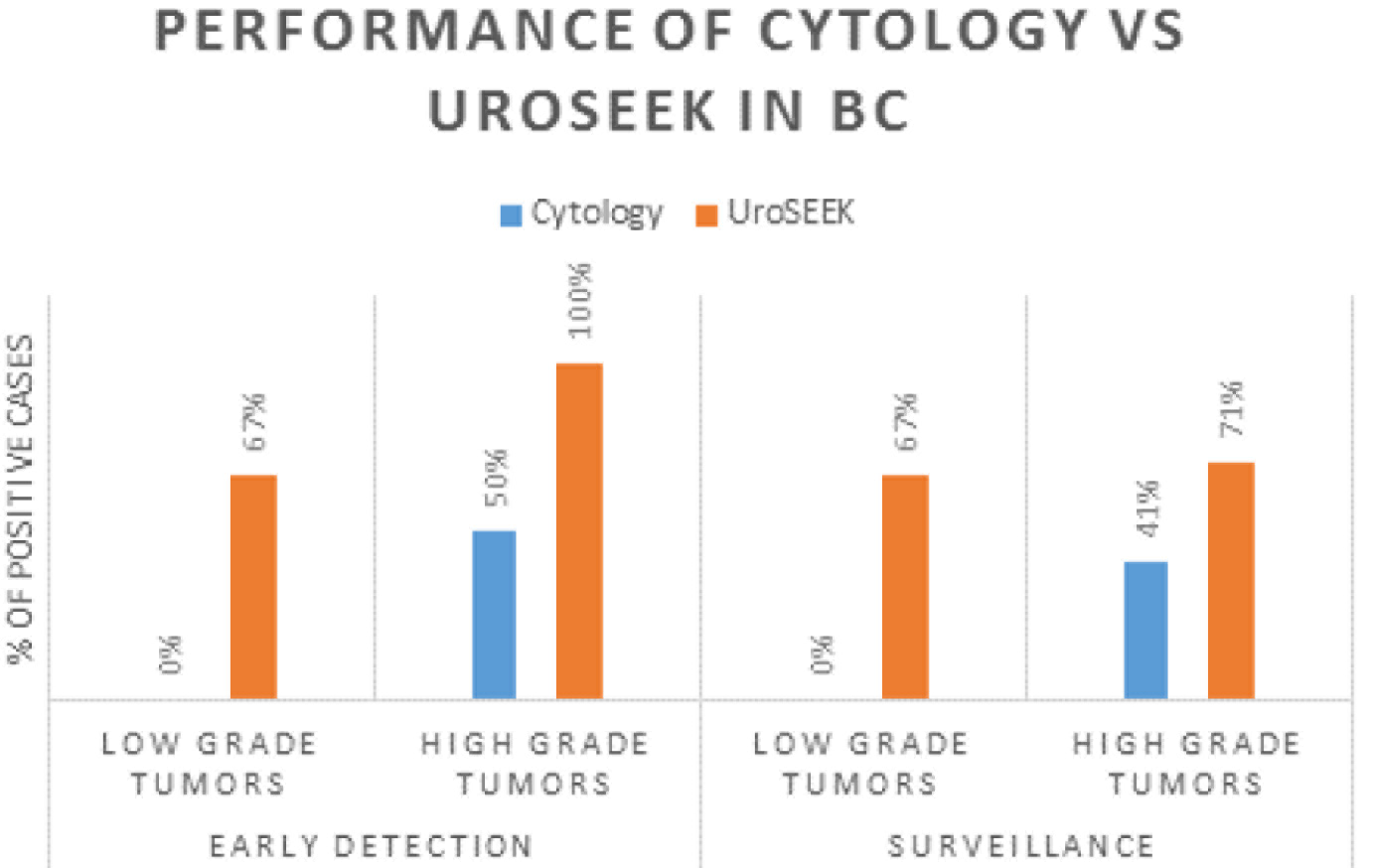
Bar graphs showing the performance of Cytology Vs. UroSEEK in diagnosis of low and high grade urothelial neoplasms in the Early Detection and Surveillance cohorts.

## Discussion

Our purpose for developing UroSEEK was not to replace cytology but rather to augment it. Cytology is a non-invasive test that is highly specific, and in expert hands nearly always indicates the presence of a BC when positive. This specificity was verified in our study: all of the 51 patients whose urine samples were positive by cytology developed biopsy-proven cancer. However, cytology is not particularly sensitive. UroSEEK adds considerably to sensitivity, as it raised the sensitivity of cytology from 43% to 95% in the Early Detection Cohort and from 25% to 71% in the Surveillance cohort. This sensitivity was highlighted by the fact that UroSEEK positive results preceded clinical diagnosis or positive cytology by months to years.

The advantage of using UroSEEK in addition to cytology was particularly evident for low-grade tumors. Cytology was negative in all 49 patients with such tumors, while 2/3 were positive with UroSEEK. Another example of the utility of the combination of UroSEEK plus cytology was evident in patients who had an equivocal cytology reading. A relatively large number of urine samples receive such an equivocal cytologic reading, even in the hands of a sub-specialized, board-certified cytopathology expert such as employed in the current study [49]. In the Early Detection Cohort, for example, 105 urine samples were scored as "atypical", and of these cases, 19% developed recurrence while the other 81% did not. UroSEEK was positive in 95% of the atypical cases that developed BC, but only in 13% of the atypical cases that did not develop cancer. These results demonstrate that UroSEEK can be used to more confidently interpret atypical cytology results.

Although UroSEEK is more sensitive than cytology, it is less specific. In this study, we were able to judge specificity in several independent ways. The first, and in some ways, most straightforward, was in a collection of urine samples from healthy individuals. In 188 such individuals, we found only one instance of a positive test, yielding a specificity of 99.5% (CI 97% to 100%). That high specificity can be considered the technical specificity of the test, but biological specificities are also important. In the Early Detection cohort, 26 of the 395 patients who did not develop BC scored positive, yielding a specificity of 93% (CI 90.50% to 96%), i.e., 6.5% false positives.

These "false positives" detected by UroSEEK could result from several factors. First, we cannot be certain that the patients whose urinary cells harbored genetic alterations did not have cancer. The follow-up period for many of patients was only one year, and cystoscopy was not generally performed. Second, it is possible that there are clonal proliferations in the bladder epithelium that increase with age. The patients in the Early Detection cohort were on average older than those in the 188 healthy individuals used as controls (40 years vs. 58 years). Though this explanation is speculative, clonal proliferations that are not considered neoplastic have been described in the bone, skin, and other tissues [50, 51]. This speculation may also explain some of the reason that mutations identified in urinary cells were not always identified in the primary tumors of the same patients. Though in the majority of cases, at least one of the mutations identified in the urine was also present in the primary tumor, this was not true in 22% of the cases in the Early Detection Cohort. In these cases, UroSEEK could be detecting clonal proliferations in the bladder epithelium that did not progress to cancer, and such proliferations may be more common in patients with BC than in the general population [50, 51]. Because only one biopsy from the primary tumor was available for comparison, it is also possible that intra-tumoral heterogeneity explains part of the discrepancies. False positives in the Surveillance Cohort could be explained in similar ways. False positives are not unique to our study; they have been observed in all other molecular assays for bladder cancer, including those that are FDA- approved [52-54]. Whether the false positives in these other assays have the same biological basis is an important area for future research.

Our study lays the conceptual and practical framework for a novel test that could inform the managements of patients with bladder cancer. Large prospective trials will be required to demonstrate the ability of UroSEEK to improve the management of patients with hematuria or dysuria or patients at risk for recurrence of BC. Before carrying out large-scale trials to evaluate such clinical utility, it is informative to predict what the performance characteristics of such a test might be. As one example, consider the use of UroSEEK plus cytology in patients presenting to their physician with microscopic hematuria or dysuria. This is a commonly encountered situation. For example, in large population-based studies involving over 80,000 individuals that participated in health screening, the fraction of individuals with micro-hematuria ranged from 2.4% to 31.1% [43, 55]. It has been estimated that 5% of such patients actually have bladder cancer [56]. In the current study, UroSEEK plus Cytology had a sensitivity of 95% and a specificity of 93% in patients of this type. This extrapolates to a positive predictive value (PPV) of 66 % (95% CI 55. to 74%) and a negative predictive value (NPV) of 99.3% (95% CI 97.3% to 99.8%). These values are well above those generally considered to be diagnostically helpful and is considerably higher than achieved in FDA-approved tests for this indication [53, 54]. The cost of a UroSEEK test is estimated to be $1000, which is comparable to that of cystoscopy, but UroSEEK is non-invasive.

## Materials and Methods

### Patients and Samples

Urine samples were collected prospectively from patients in four participating institutions including Johns Hopkins Hospital, Baltimore, MD, USA; A.C. Camargo Cancer Center, Sao Paulo, Brazil; Osaka University Hospital, Osaka, Japan; and Hacettepe University Hospital, Ankara, Turkey. The study was approved by the institutional Review Boards of Johns Hopkins Hospital and all other participating institutions. Proper material transfer agreements were obtained. Patients with a known history of malignancy other than bladder cancer were excluded from the study. The study included two cohorts of patients. The Early Detection cohort comprised patients who were referred to a urology clinic in one of the above hospitals because of hematuria or lower urinary tract symptoms (Supplementary File 1) (570 Patients). The second cohort (322 patients) represented patients with prior established diagnosis of BC who are on surveillance for disease recurrence (Surveillance Cohort). As noted in the main text, these patients’ primary tumors harbored mutations in at least one of the 11 genes assessed through the multiplex or singleplex assays. A minimum follow-up of 12 months was from date of urine collection was required for cases with no evidence of incident or recurrent tumors in the Early Detection or Surveillance cohorts, respectively. Urine samples were collected prior to any procedures, such as cystoscopy, performed during the patients’ visits. A total of 892 urine samples were analyzed in the study, composed of two type of samples. The first was residual urinary cells after processing with standard BD SurePath™ liquid-based cytology protocols (Becton Dickinson and Company; Franklin Lakes, NJ, USA). To allow for standard-of-care, residual SurePath^®^ fluids were kept refrigerated for 6-8 weeks prior to submission for DNA purification to allow for any potential need for repeat cytology processing of the same sample. The second sample type was composed of bio-banked fresh urine samples in which 15 - 25 mL of voided urine samples were stored at 4°C for up to 60 min prior to centrifugation (10 min at 500 g) and the pellets stored at minus 80°C prior to DNA purification. Urines from 188 healthy individuals of average age 26 were also obtained and processed identically to the bio-banked fresh urine samples.

Formalin-fixed paraffin-embedded (FFPE) tumor tissue samples from trans-urethral resections (TURB) or cystectomies were collected in 413 of the 892 cases. When several different tumors from the same patient were available (because of recurrences), the earliest tumor tissue obtained following the donation of the urine sample was used in the Early Detection Cohort. In the surveillance cohort, the tumors preceding the donation of the urine sample was used in 146 of the 322 patients. In the other 176 Surveillance cases, the earliest tissue obtained following the donation of the urine sample was used. A genitourinary pathologist reviewed all histologic slides to confirm the diagnosis and select a representative tumor area with as high tumor cellularity as possible for that case. Corresponding FFPE blocks were cored with a sterile 16-gauge needle. One to three cores were obtained per tumor and placed in 1.5-mL sterile tubes for DNA purification, as previously described [14]. Electronic medical records were reviewed to obtain medical history and follow up data in all patients.

### Mutation analysis

Three separate assays were used to search for abnormalities in urinary cell DNA. First, a multiplex PCR was used to detect mutations in regions of ten genes commonly mutated in urologic malignancies *CDKN2A, ERBB2, FGFR3, HRAS, KRAS, MET, MLL, PIK3CA, TP53, and VHL* [33-40]. The 57 primer pairs used for this multiplex PCR were divided in a total of three multiplex reactions, each containing non-overlapping amplicons (Supplementary Table 13) These primers were used to amplify DNA in 25 uL reactions as previously described [42] except that 15 cycles was used for the initial amplification. Second, the *TERT* promoter region was evaluated. A single amplification primer was used to amplify a 73-bp segment containing the region of the *TERT* promoter known to harbor mutations in BC [14]. The conditions used to amplify it were the same as used in the multiplex reactions described above except that Phusion GC Buffer (Thermo-Fisher) instead of HF buffer was used and 20 cycles were used for the initial amplification. Note that the *TERT* promoter region could not be included in the multiplex PCR because of the high GC content of the former. PCR products were purified with AMPure XP beads (Beckman Coulter, PA, USA) and 0.25% of the purified PCR products (multiplex) or 0.0125% of the PCR products (*TERT* singleplex) were then amplified in a second round of PCR, as described in [57]. The PCR products from the second round of amplification were then purified with AMPure and sequenced on an Illumina instrument. For each mutation identified, the mutant allele frequency (MAF) was determined by dividing the number of uniquely identified reads with mutations [42] by the number of total uniquely identified reads. Each DNA sample was assessed in two independent PCRs, for both the *TERT* promoter and multiplex assays, and samples were scored as positive only if both PCRs showed the same mutation. The mutant allele frequencies and number of UIDs listed in the Supplementary Tables refer to the average of the two independent assays.

To evaluate the statistical significance of putative mutations, we assessed DNA from white blood cells of 188 unrelated normal individuals. A variant observed in the samples from cancer patient was only scored as a mutation if it was observed at a much higher MAF than observed in normal WBCs. Specifically, the classification of a sample’s ctDNA status was based on two complementary criteria applied to each mutation: 1) the difference between the average MAF in the sample of interest and the corresponding maximum MAF observed for that same mutation in a set of controls, and 2) the Stouffer’s Z-score obtained by comparing the MAF in the sample of interest to a distribution of normal controls. To calculate the Z-score, the MAF in the sample of interest was first normalized based on the mutation-specific distributions of MAFs observed among all controls. Following this mutation-specific normalization, a P-value was obtained by comparing the MAF of each mutation in each well with a reference distribution of MAFs built from normal controls where all mutations were included. The Stouffer’s Z-score was then calculated from the p-values of two wells, weighted by their number of UIDs. The sample was classified as positive if either the difference or the Stouffer’s Z-score of its mutations was above the thresholds determined from the normal WBCs. The threshold for the difference parameter was defined by the highest MAF observed in any normal WBCs. The threshold for the Stouffer’s Z-score was chosen to allow one false positive among the 188 normal urine samples studied.

#### Analysis of aneuploidy

Aneuploidy was assessed with FastSeqS, which uses a single primer pair to amplify ~38,000 loci scattered throughout the genome [41]. After massively parallel sequencing, gains or losses of each of the 39 chromosome arms covered by the assay were determined using a bespoke statistical learning method described elsewhere (Deauville et al., in preperation). A Support vector machine (SVM) was used to discriminate between aneuploid and euploid samples. The SVM was trained using 3150 low neoplastic cell fraction synthetic aneuploid samples and 677 euploid peripheral white blood cell (WBC) samples. Samples were scored as positive when the genome-wide aneuploidy score was >0.7 and there was at least one gain or loss of a chromosome arm.

#### Identity checks

A multiplex reaction containing 26 primers detecting 31 common SNPs on chromosomes 10 and 20 was performed using the amplification conditions described 638 above for the multiplex PCR. The 26 primers used for this identity evaluation are listed in Supplementary Table 14.

### Statistical Analysis

Performance characteristics of urine cytology, UroSEEK and its three components was calculated using using MedCalc statistical software, online version (https://www.medcalc.org/calc/diagnostic_test.php).

## Acknowledgments

The generous support provided by Henry and Marsha Laufer is gratefully acknowledged, as is support from the The Virginia and D.K. Ludwig Fund for Cancer Research, The Commonwealth Foundation, and The Conrad R. Hilton Foundation. All sequencing was performed at the Sol Goldman Sequencing Facility at Johns Hopkins. This work was also supported by grants from the NIH (Grants CA-77598, CA 06973, and GM 07309). The authors appreciate the medical illustrations skillfully designed by Kathleen Gebhart (Media Services, Stony Brook Univ.).

## Author contributions

Simeon Springer: Acquisition of data, Analysis and interpretation of data, Drafting and revising the article

Maria Del Carmen Rodriguez Pena: Acquisition of data, Analysis and interpretation of data, specimen collection, Drafting and revising the article

Lu Li, Acquisition of data, Analysis and interpretation of data

Chris Douville, Acquisition of data, Analysis and interpretation of data

Yuxuan Wang: Acquisition of data, Analysis and interpretation of data, Drafting and revising the article

Josh Cohen, Acquisition of data, Analysis and interpretation of data

Diana Taheri, Acquisition of data, Analysis and interpretation of data, specimen collection

Natalie Silliman, Acquisition of data, Analysis and interpretation of data

Joy Schaeffer, Acquisition of data, Analysis and interpretation of data

Janine Ptak, Acquisition of data, Analysis and interpretation of data

Lisa Dobbyn, Acquisition of data, Analysis and interpretation of data

Maria Papoli, Acquisition of data, Analysis and interpretation of data

Isaac Kinde, Conception and design

Bahman Afsari, Analysis and interpretation of data

Aline C. Tregnago: Acquisition of data, Analysis and interpretation of data, specimen collection

Stephania M. Bezerra, Specimen contribution, Acquisition of data

Christopher VandenBussche: Specimen contribution, Revising the article

Kazutoshi Fujita: Specimen contribution, Acquisition of data

Dilek Ertoy: Specimen contribution, Acquisition of data

Isabela W. Cunha: Specimen contribution, Acquisition of data

Lijia Yu, Analysis and interpretation of data

Mark Schoenberg, Specimen contribution

Trinity J. Bivalacqua, Specimen contribution

Kathleen G. Dickman, Revising the article

Arthur P. Grollman, Revising the article

Luis A Diaz, Conception and design, Revising the article

Rachel Karchin, Design of aneuploidy caller, Analysis and interpretation of data.

Ralph Hruban Conception and design

Cristian Tomasetti, Design of algorithm for sequencing data analysis, Analysis and interpretation of data.

Nickolas Papadopoulos: Conception and design, Acquisition of data, Analysis and interpretation of data, Drafting and revising the article, Specimen contribution

Kenneth W. Kinzler: Conception and design, Acquisition of data, Analysis and interpretation of data, Drafting and revising the article, Specimen contribution

Bert Vogelstein: Conception and design, Acquisition of data, Analysis and interpretation of data, Drafting and revising the article, Specimen contribution

George J. Netto: Conception and design, Acquisition of data, Analysis and interpretation of data, Drafting and revising the article, Specimen contribution

## References

1. Siegel RL, Miller KD, Jemal A (2017) Cancer Statistics, 2017. CA Cancer J Clin 67:7–30

2. Netto GJ (2013) Clinical applications of recent molecular advances in urologic malignancies: no longer chasing a “mirage”?. Adv Anat Pathol 20:175–203

3. Netto GJ, Tafe LJ (2016) Emerging Bladder Cancer Biomarkers and Targets of Therapy. Urol Clin North Am 43:63–76

4. Lotan Y, Roehrborn CG (2003) Sensitivity and specificity of commonly available bladder tumor markers versus cytology: results of a comprehensive literature review and meta-analyses. Urology 61:109–18; discussion 118

5. Zhang ML, Rosenthal DL, VandenBussche CJ (2016) The cytomorphological features of low-grade urothelial neoplasms vary by specimen type. Cancer Cytopathol 124:552–564

6. Kawauchi S, Sakai H, Ikemoto K, Eguchi S, Nakao M, Takihara H, Shimabukuro T, Furuya T, Oga A, Matsuyama H, Takahashi M, Sasaki K (2009) 9p21 Index as Estimated by Dual-Color Fluorescence in Situ Hybridization is Useful to Predict Urothelial Carcinoma Recurrence in Bladder Washing Cytology. Hum Pathol 40:1783–1789

7. Kruger S, Mess F, Bohle A, Feller AC (2003) Numerical aberrations of chromosome 17 and the 9p21 locus are independent predictors of tumor recurrence in non-invasive transitional cell carcinoma of the urinary bladder. Int J Oncol 23:41–48

8. Skacel M, Fahmy M, Brainard JA, Pettay JD, Biscotti CV, Liou LS, Procop GW, Jones JS, Ulchaker J, Zippe CD, Tubbs RR (2003) Multitarget fluorescence in situ hybridization assay detects transitional cell carcinoma in the majority of patients with bladder cancer and atypical or negative urine cytology. J Urol 169:2101–2105

9. Sarosdy MF, Kahn PR, Ziffer MD, Love WR, Barkin J, Abara EO, Jansz K, Bridge JA, Johansson SL, Persons DL, Gibson JS (2006) Use of a multitarget fluorescence in situ hybridization assay to diagnose bladder cancer in patients with hematuria. J Urol 176:44–47

10. Moonen PM, Merkx GF, Peelen P, Karthaus HF, Smeets DF, Witjes JA (2007) UroVysion compared with cytology and quantitative cytology in the surveillance of non-muscle-invasive bladder cancer. Eur Urol 51:1275–80; discussion 1280

11. Fradet Y, Lockhard C (1997) Performance characteristics of a new monoclonal antibody test for bladder cancer: ImmunoCyt trade mark. Can J Urol 4:400–405

12. Yafi FA, Brimo F, Steinberg J, Aprikian AG, Tanguay S, Kassouf W (2015) Prospective analysis of sensitivity and specificity of urinary cytology and other urinary biomarkers for bladder cancer. Urol Oncol 33:66.e25–66.e31

13. Serizawa RR, Ralfkiaer U, Steven K, Lam GW, Schmiedel S, Schuz J, Hansen AB, Horn T, Guldberg P (2010) Integrated genetic and epigenetic analysis of bladder cancer reveals an additive diagnostic value of FGFR3 mutations and hypermethylation events. Int J Cancer

14. Kinde I, Munari E, Faraj SF, Hruban RH, Schoenberg M, Bivalacqua T, Allaf M, Springer S, Wang Y, Diaz LA,Jr, Kinzler KW, Vogelstein B, Papadopoulos N, Netto GJ (2013) TERT promoter mutations occur early in urothelial neoplasia and are biomarkers of early disease and disease recurrence in urine. Cancer Res 73:7162–7167

15. Hurst CD, Platt FM, Knowles MA (2014) Comprehensive mutation analysis of the TERT promoter in bladder cancer and detection of mutations in voided urine. Eur Urol 65:367–369

16. Wang K, Liu T, Ge N, Liu L, Yuan X, Liu J, Kong F, Wang C, Ren H, Yan K, Hu S, Xu Z, Bjorkholm M, Fan Y, Zhao S, Liu C, Xu D (2014) TERT promoter mutations are associated with distant metastases in upper tract urothelial carcinomas and serve as urinary biomarkers detected by a sensitive castPCR. Oncotarget 5:12428–12439

17. Ralla B, Stephan C, Meller S, Dietrich D, Kristiansen G, Jung K (2014) Nucleic acid-based biomarkers in body fluids of patients with urologic malignancies. Crit Rev Clin Lab Sci 51:200–231

18. Ellinger J, Muller SC, Dietrich D (2015) Epigenetic biomarkers in the blood of patients with urological malignancies. Expert Rev Mol Diagn 15:505–516

19. Bansal N, Gupta A, Sankhwar SN, Mahdi AA (2014) Low- and high-grade bladder cancer appraisal via serum-based proteomics approach. Clin Chim Acta 436:97–103

20. Goodison S, Chang M, Dai Y, Urquidi V, Rosser CJ (2012) A multi-analyte assay for the non-invasive detection of bladder cancer. PLoS One 7:e47469

21. Allory Y, Beukers W, Sagrera A, Flandez M, Marques M, Marquez M, van der Keur KA, Dyrskjot L, Lurkin I, Vermeij M, Carrato A, Lloreta J, Lorente JA, Carrillo-de Santa Pau E, Masius RG, Kogevinas M, Steyerberg EW, van Tilborg AA, Abas C, Orntoft TF, Zuiverloon TC, Malats N, Zwarthoff EC, Real FX (2014) Telomerase reverse transcriptase promoter mutations in bladder cancer: high frequency across stages, detection in urine, and lack of association with outcome. Eur Urol 65:360–366

22. Huang FW, Hodis E, Xu MJ, Kryukov GV, Chin L, Garraway LA (2013) Highly recurrent TERT promoter mutations in human melanoma. Science 339:957–959

23. Killela PJ, Reitman ZJ, Jiao Y, Bettegowda C, Agrawal N, Diaz LA,Jr, Friedman AH, Friedman H, Gallia GL, Giovanella BC, Grollman AP, He TC, He Y, Hruban RH, Jallo GI, Mandahl N, Meeker AK, Mertens F, Netto GJ, Rasheed BA, Riggins GJ, Rosenquist TA, Schiffman M, Shih I, Theodorescu D, Torbenson MS, Velculescu VE, Wang TL, Wentzensen N, Wood LD, Zhang M, McLendon RE, Bigner DD, Kinzler KW, Vogelstein B, Papadopoulos N, Yan H (2013) TERT promoter mutations occur frequently in gliomas and a subset of tumors derived from cells with low rates of self-renewal. Proc Natl Acad Sci U S A 110:6021–6026

24. Scott GA, Laughlin TS, Rothberg PG (2014) Mutations of the TERT promoter are common in basal cell carcinoma and squamous cell carcinoma. Mod Pathol 27:516–523

25. Horn S, Figl A, Rachakonda PS, Fischer C, Sucker A, Gast A, Kadel S, Moll I, Nagore E, Hemminki K, Schadendorf D, Kumar R (2013) TERT promoter mutations in familial and sporadic melanoma. Science 339:959–961

26. Allory Y, Beukers W, Sagrera A, Flandez M, Marques M, Marquez M, van der Keur KA, Dyrskjot L, Lurkin I, Vermeij M, Carrato A, Lloreta J, Lorente JA, Carrillo-de Santa Pau E, Masius RG, Kogevinas M, Steyerberg EW, van Tilborg AA, Abas C, Orntoft TF, Zuiverloon TC, Malats N, Zwarthoff EC, Real FX (2014) Telomerase reverse transcriptase promoter mutations in bladder cancer: high frequency across stages, detection in urine, and lack of association with outcome. Eur Urol 65:360–366

27. Cowan ML, Springer S, Nguyen D, Taheri D, Guner G, Mendoza Rodriguez MA, Wang Y, Kinde I, Del Carmen Rodriguez Pena M, VandenBussche CJ, Olson MT, Cunha I, Fujita K, Ertoy D, Kinzler K, Bivalacqua T, Papadopoulos N, Vogelstein B, Netto GJ (2016) Detection of TERT promoter mutations in primary adenocarcinoma of the urinary bladder. Hum Pathol 53:8–13

28. Nguyen D, Taheri D, Springer S, Cowan M, Guner G, Mendoza Rodriguez MA, Wang Y, Kinde I, VandenBussche CJ, Olson MT, Ricardo BF, Cunha I, Fujita K, Ertoy D, Kinzler KW, Bivalacqua TJ, Papadopoulos N, Vogelstein B, Netto GJ (2016) High prevalence of TERT promoter mutations in micropapillary urothelial carcinoma. Virchows Arch 469:427–434

29. Rodriguez Pena MDC, Tregnago AC, Eich ML, Springer S, Wang Y, Taheri D, Ertoy D, Fujita K, Bezerra SM, Cunha IW, Raspollini MR, Yu L, Bivalacqua TJ, Papadopoulos N, Kinzler KW, Vogelstein B, Netto GJ (2017) Spectrum of genetic mutations in de novo PUNLMP of the urinary bladder. Virchows Arch

30. Cheng L, Montironi R, Lopez-Beltran A (2017) TERT Promoter Mutations Occur Frequently in Urothelial Papilloma and Papillary Urothelial Neoplasm of Low Malignant Potential. Eur Urol 71:497–498

31. International Agency for Research on Cancer. (2016) WHO Classification of Tumours of the Urinary System and Male Genital Organs. World Health Organization; 4 edition

32. Netto GJ (2011) Molecular biomarkers in urothelial carcinoma of the bladder: are we there yet?. Nat Rev Urol 9:41–51

33. Netto GJ (2011) Molecular biomarkers in urothelial carcinoma of the bladder: are we there yet?. Nat Rev Urol 9:41–51

34. Mo L, Zheng X, Huang HY, Shapiro E, Lepor H, Cordon-Cardo C, Sun TT, Wu XR (2007) Hyperactivation of Ha-ras oncogene, but not Ink4a/Arf deficiency, triggers bladder tumorigenesis. J Clin Invest 117:314–325

35. Sarkis AS, Dalbagni G, Cordon-Cardo C, Zhang ZF, Sheinfeld J, Fair WR, Herr HW, Reuter VE (1993) Nuclear overexpression of p53 protein in transitional cell bladder carcinoma: a marker for disease progression. J Natl Cancer Inst 85:53–59

36. Lin HH, Ke HL, Huang SP, Wu WJ, Chen YK, Chang LL (2010) Increase sensitivity in detecting superficial, low grade bladder cancer by combination analysis of hypermethylation of E-cadherin, p16, p14, RASSF1A genes in urine. Urol Oncol 28:597–602

37. Sarkis AS, Dalbagni G, Cordon-Cardo C, Melamed J, Zhang ZF, Sheinfeld J, Fair WR, Herr HW, Reuter VE (1994) Association of P53 nuclear overexpression and tumor progression in carcinoma in situ of the bladder. J Urol 152:388–392

38. Sarkis AS, Bajorin DF, Reuter VE, Herr HW, Netto G, Zhang ZF, Schultz PK, Cordon-Cardo C, Scher HI (1995) Prognostic value of p53 nuclear overexpression in patients with invasive bladder cancer treated with neoadjuvant MVAC. J Clin Oncol 13:1384–1390

39. Wu XR (2005) Urothelial tumorigenesis: a tale of divergent pathways. Nat Rev Cancer 5:713–725

40. Cancer Genome Atlas Research Network (2014) Comprehensive molecular characterization of urothelial bladder carcinoma. Nature 507:315–322

41. Kinde I, Papadopoulos N, Kinzler KW, Vogelstein B (2012) FAST-SeqS: a simple and efficient method for the detection of aneuploidy by massively parallel sequencing. PLoS One 7:e41162

42. Kinde I, Wu J, Papadopoulos N, Kinzler KW, Vogelstein B (2011) Detection and quantification of rare mutations with massively parallel sequencing. Proc Natl Acad Sci U S A 108:9530–9535

43. Wein AJ, Kavoussi LR, Novick AC, Partin AW, Peters CA (2012) Campbell-Walsh Urology. Saunders, Philadelphia

44. Mishriki SF, Nabi G, Cohen NP (2008) Diagnosis of urologic malignancies in patients with asymptomatic dipstick hematuria: prospective study with 13 years’ follow-up. Urology 71:13–16

45. Netto GJ, Epstein JI (2010) Theranostic and prognostic biomarkers: genomic pplications in urological malignancies. Pathology 42:384–394

46. Cancer Genome Atlas Research Network (2014) Comprehensive molecular characterization of urothelial bladder carcinoma. Nature 507:315–322

47. Stern JL, Theodorescu D, Vogelstein B, Papadopoulos N, Cech TR (2015) Mutation of the TERT promoter, switch to active chromatin, and monoallelic TERT expression in multiple cancers. Genes Dev 29:2219–2224

48. Vogelstein B, Papadopoulos N, Velculescu VE, Zhou S, Diaz LA,Jr, Kinzler KW (2013) Cancer genome landscapes. Science 339:1546–1558

49. Barkan GA, Wojcik EM, Nayar R, Savic-Prince S, Quek ML, Kurtycz DF, Rosenthal DL (2016) The Paris System for Reporting Urinary Cytology: The Quest to Develop a Standardized Terminology. Adv Anat Pathol 23:193–201

50. Chai H, Brown RE (2009) Field effect in cancer-an update. Ann Clin Lab Sci 39:331–337

51. Takahashi T, Habuchi T, Kakehi Y, Mitsumori K, Akao T, Terachi T, Yoshida O (1998) Clonal and chronological genetic analysis of multifocal cancers of the bladder and upper urinary tract. Cancer Res 58:5835–5841

52. Gopalakrishna A, Fantony JJ, Longo TA, Owusu R, Foo WC, Dash R, Denton BT, Inman BA (2017) Anticipatory Positive Urine Tests for Bladder Cancer. Ann Surg Oncol 24:1747–1753

53. Hajdinjak T (2008) UroVysion FISH test for detecting urothelial cancers: meta-analysis of diagnostic accuracy and comparison with urinary cytology testing. Urol Oncol 26:646–651

54. Dimashkieh H, Wolff DJ, Smith TM, Houser PM, Nietert PJ, Yang J (2013) Evaluation of urovysion and cytology for bladder cancer detection: a study of 1835 paired urine samples with clinical and histologic correlation. Cancer Cytopathol 121:591–597

55. Davis R, Jones JS, Barocas DA, Castle EP, Lang EK, Leveillee RJ, Messing EM, Miller SD, Peterson AC, Turk TM, Weitzel W, American Urological Association (2012) Diagnosis, evaluation and follow-up of asymptomatic microhematuria (AMH) in adults: AUA guideline. J Urol 188:2473–2481

56. Khadra MH, Pickard RS, Charlton M, Powell PH, Neal DE (2000) A prospective analysis of 1,930 patients with hematuria to evaluate current diagnostic practice. J Urol 163:524–527

57. Wang Y, Sundfeldt K, Mateoiu C, Shih I, Kurman RJ, Schaefer J, Silliman N, Kinde I, Springer S, Foote M, Kristjansdottir B, James N, Kinzler KW, Papadopoulos N, Diaz LA, Vogelstein B (2016) Diagnostic potential of tumor DNA from ovarian cyst fluid. Elife 5:10.7554/eLife.15175

